# The ventral tegmental area dopamine to lateral amygdala projection supports cocaine cue associative learning

**DOI:** 10.1101/2023.08.22.554187

**Authors:** Dana M. Smith, Mary M. Torregrossa

**Affiliations:** Department of Psychiatry, University of Pittsburgh, 450 Technology Drive, Pittsburgh, PA 15219, USA; Center for Neuroscience, University of Pittsburgh, 4200 Fifth Ave, Pittsburgh, PA 15213, USA

## Abstract

Learning and memory mechanisms are critically involved in drug craving and relapse. Environmental cues paired with repeated drug use acquire incentive value such that exposure to the cues alone can trigger craving and relapse. The amygdala, particularly the lateral amygdala (LA), underlies cue-related learning processes that assign valence to environmental stimuli including drug-paired cues. Evidence suggests that the ventral tegmental area (VTA) dopamine (DA) projection to the LA participates in encoding reinforcing effects that act as a US in conditioned cue reward-seeking as DA released in the amygdala is important for emotional and behavioral functions. Here we used chemogenetics to manipulate these VTA DA inputs to the LA to determine the role of this projection for acquisition of drug-cue associations and reinstatement of drug-seeking. We found inhibiting DA input to the LA during cocaine self-administration slowed acquisition and weakened the ability of the previously cocaine-paired cue to elicit cocaine-seeking. Conversely, exciting the projection during self-administration boosted the salience of the cocaine-paired cue as indicated by enhanced responding during cue-induced reinstatement. Importantly, interfering with DA input to the LA had no impact on the ability of cocaine to elicit a place preference or induce reinstatement in response to a priming cocaine injection. Overall, we show that manipulation of projections underlying DA signaling in the LA may be useful for developing therapeutic interventions for substance use disorders.

## Introduction

Learning and memory mechanisms are critically involved in drug craving and relapse^1^. As environmental cues become paired with repeated drug use, these cues acquire incentive and motivational value such that exposure to the cues alone can trigger craving and relapse. The amygdala is a key region underlying cue-related learning processes that assign valence to environmental stimuli including drug-paired cues^2^. Many associative learning studies have focused on threat processing and fear conditioning. In this work, an auditory conditioned stimulus (CS) is paired with an aversive footshock unconditioned stimulus (US), leading to threat appropriate responses occurring in response to the CS alone. Experiments revealed that the lateral amygdala (LA) mediates this learning via integration of sensory inputs from auditory thalamus and cortex^3–7^ with the negative footshock effects via projections from somatosensory regions^8^.

The traditional model for this process is that an initially weak afferent pathway carrying sensory information about the CS and strong afferents carrying US information converge in the LA. Through plasticity mechanisms, excitatory synapses carrying CS information are strengthened^7,9,10^. In the context of drugs of abuse, drug-cue associations also develop through enhancing CS inputs from auditory thalamus onto LA principal neurons^6^. Yet, the regions providing inputs to the LA to encode the reinforcing effects of cocaine that serve as a US are not well characterized.

Evidence suggests that the ventral tegmental area (VTA) projection to the LA participates in encoding reinforcing effects of drugs that act as a US in conditioned cue drug-seeking as DA activity in the LA has been shown to be necessary for maintenance of drug-cue associations and conditioned reinstatement of drug-seeking^11–14^. Much of the DA in LA originates from the VTA and likely modulates learning through altering the excitability of amygdala projection neurons and interneurons^15^. These VTA DA to LA projections contribute to aversive and appetitive stimulus-outcome learning^16,17^ and stimulating DA terminals in the amygdala has been shown to strengthen the motivational impact of previously learned reward cues^18^. Here, we used various chemogenetic approaches to manipulate VTA DA to LA projections during cocaine self-administration and subsequent reinstatement to determine their role in cocaine cue associative learning. Our results extend the current knowledge of the role of DA in the LA to show an important contribution of these projections in associative learning and point to a relevant pathway in the development of substance use disorder.

## Methods

### Animals

Adult male and female Sprague Dawley rats (Envigo) were pair-housed in auto-ventilated racks with automated watering in a temperature-and humidity-controlled room and maintained on a 12h light/dark cycle. Following catheter implantation, rats were single housed and food restricted 24hr before the start of training (maintained at 95% free-feeding weight). In experiments using TH-Cre rats, both sexes were bred in house from Tyrosine Hydroxylase TH-Cre knockin (HsdSage:SD-TH^em1(IRES-CRE)Sage^) dams and wild-type males or the offspring of these breeding pairs. Genotypes were determined by Transnetyx® and heterozygous TH-Cre animals were used. All procedures were conducted in accordance with the National Institutes of Health’s *Guide for Care and Use of Laboratory Animals* and approved by the University of Pittsburgh’s Institutional Animal Care and Use Committee.

### Viral constructs

For chemogenetics in TH-Cre rats, the LA (males in mm from bregma, AP: -3.0, ML: ±5.0, DV: -7.9 mm; females, AP: -2.8, ML: ±4.8; DV: -7.8 mm) was injected bilaterally (1μl/hemisphere) with retrograde adeno-associated virus containing a double-floxed, inverted open reading frame (DIO) sequence for the mCherry-tagged hM4Di or mCherry control (hSyn-DIO-hM4D(Gi)-mCherry, hSyn-DIO-mCherry, Addgene). A custom AAV2retro.PRSx8.HA-hM4D.SV40 targeting noradrenergic neurons (Penn Vector Core, kindly provided by Dr. Gary Aston-Jones) was bilaterally infused into the LA of a separate group of Sprague Dawley rats. In other chemogenetic experiments, the VTA (both sexes: AP: -5.5, ML: ± 0.9, DV: -8.20) was injected bilaterally (1μl/hemisphere) with AAV2 containing the mCherry-tagged hM4Di or hM3Dq DREADDs or EGFP (CaMKIIα - hM4D(Gi)-mCherry, CaMKIIα-hM3D(Gq)-mCherry, CaMKIIα-EGFP, Addgene). See Supplement for delivery details.

### Drugs

Cocaine hydrochloride (provided by NIDA) was dissolved (intravenous self-administration: 2mg/mL; intraperitoneal injections: 5mg/mL) in 0.9% sterile saline (ThermoFisher) and filter-sterilized. Clozapine-N-oxide (CNO, provided by NIDA) was dissolved in 5% DMSO in 0.9% sterile saline (1 or 3mg/mL) and injected intraperitoneally 30 min before behavior. For intracranial injection, CNO was dissolved in 0.5% DMSO in artificial CSF (ACSF, Fisher Scientific; 1mM/0.3 µl; total dose, ∼100 ng/hemisphere) and microinjected 5 min before behavior as previously described^18^.

### Self-administration

Rats were implanted with intravenous catheters (see Supplement) and trained to self-administer cocaine (2mg/mL) in daily 1hr sessions on a FR1 schedule with 10s timeout. Sessions began with house light illumination and lever insertion. An active lever press (side counterbalanced between animals) produced an infusion paired with a 10s audiovisual cue (tone and cue light illumination above active lever). Inactive lever presses were recorded, but had no consequence. Pump durations were adjusted daily according to body weight to deliver 1mg/kg/infusion. Sessions terminated after 1hr or 30 infusions. Rats received systemic CNO (1mg/kg) or vehicle injection (i.p.) 30 min prior to the session. Rats implanted with intracranial LA cannula (see Supplement) received CNO or ACSF microinfusions 5 min before each session. Rats receiving microinfusions were restricted to 5 days of self-administration and first trained on sucrose pellet self-administration in a 6hr session on a FR1 schedule.

### Instrumental Extinction

For 7-14 days, during daily 1hr sessions, lever responses were recorded but had no consequences. Extinction continued until rats made an average of <25 active presses on the last 2 extinction days.

### Cue-induced reinstatement

During a 1hr session, an active press resulted in a 10s presentation of the cocaine-associated cue on an FR1 schedule. Inactive presses were recorded but had no consequences. For repeated reinstatement, rats received i.p. injections of vehicle, 1mg/kg, or 3mg/kg CNO with the order counterbalanced between animals. Test days were separated by ∼2 days of instrumental re-extinction.

### Cocaine primed reinstatement

Immediately prior to a 1hr session, rats received a 10mg/kg cocaine i.p. injection. Rats had access to levers and presses were recorded, but had no consequences. Repeated reinstatement involved i.p. injections of vehicle, 1mg/kg, or 3mg/kg CNO with the order counterbalanced between rats and each test was separated by ∼2 days of re-extinction.

### Conditioned Place Preference (CPP)

Rats were tested in a custom CPP apparatus (see Supplement). At baseline, rats were placed in the center of the apparatus to explore for 15 min with time spent in each chamber scored. Animals were considered in a chamber if 4 paws were located inside, while 2 paws outside was an exit. Following baseline, rats had 6 conditioning sessions. On alternate days, rats were injected (i.p.) with cocaine and restricted to a randomly assigned (paired) compartment for 20 min or injected with saline and restricted to the unpaired compartment for 20 min. Males received 20 mg/kg cocaine and females received 5 mg/kg cocaine, as these doses have been shown to produce equivalent levels of CPP between the sexes^19^. Any injection (i.p.) of 1mg/kg CNO or vehicle occurred 30 min prior to the conditioning session, while LA CNO or vehicle microinfusions occurred 5 min before the session. Animals received the same treatment each conditioning day based on their group assignment. The 15 min preference test was conducted 24hr after the final conditioning session and the divider separating the chambers was removed. Rats were placed in the center and time spent in each of the chambers was scored.

### Statistics

Behavioral data were collected using MedPC software and experimenters were blinded to the rats’ treatment conditions. Statistical analyses were performed using GraphPad Prism and SPSS Statistics Software with significance set at p<0.05.

## Results

### Chemogenetic inhibition of VTA DA input to LA slows acquisition of cocaine self-administration

To determine the contribution of VTA DA input to LA in cocaine cue associative learning, we inhibited the projection as animals learned self-administration. Prior to self-administration, TH-Cre rats received bilateral LA injections of a Cre-dependent retrograde AAV expressing hM4Di (Gi-DREADD) or mCherry (control). Rats were then trained to self-administer cocaine (1mg/kg/infusion) for 14 days while receiving injections (i.p) of 1 mg/kg CNO (Gi-DREADD and mCherry control group) or vehicle (Gi-DREADD group) before each session (Figure 1A). An ANOVA with within-subjects factor of training day and between-subjects factors of treatment group and sex was performed. There was no main effect of sex on infusions earned as both males (n=9) and females (n=9) earned a similar number of infusions (F_(1,12)_=0.33, p=0.58). There were no sex differences in active (F_(1,12)_=0.41, p=0.54) or inactive presses (F_(1,12)_=0.55, p=0.47). Some of the individual experimental groups were small (n=5) with 2-3 animals of each sex per group and were not powered to detect sex differences. Given the overall lack of sex differences in the majority of measures, data were analyzed with both sexes combined.

**Figure 1.**
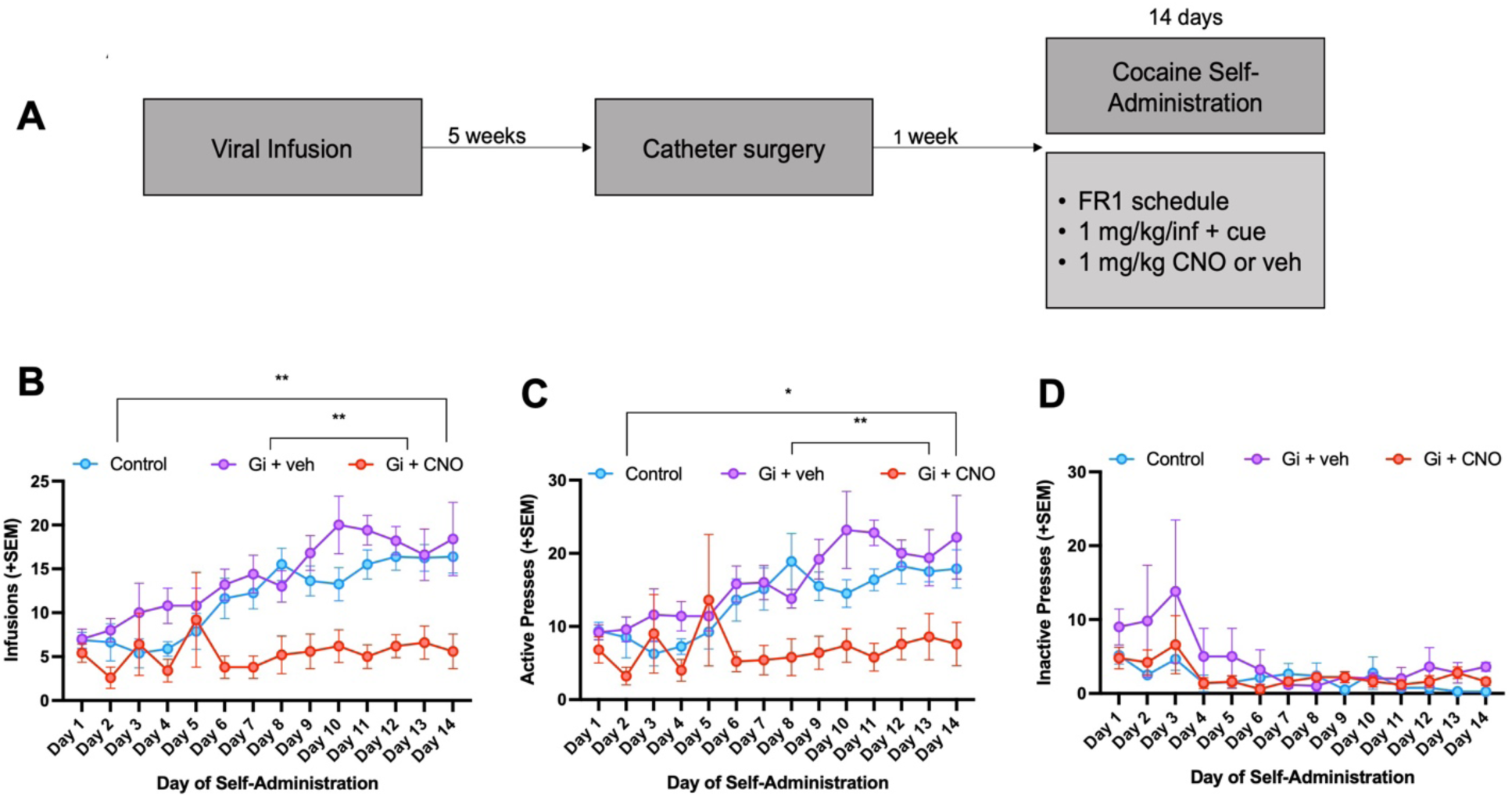
Inhibition of VTA DA input to LA slows acquisition of cocaine self-administration. Rats (n=18, Control=8, Gi + veh=5, Gi + CNO=5) self-administered cocaine (1mg/kg/infusion) on a FR1 schedule and received i.p. injections of either 1mg/kg CNO or vehicle prior to each session. Experimental timeline (**A**). Gi + CNO rats earned fewer cocaine infusions than Gi + veh and control rats (**B**). Gi + CNO rats made fewer active presses than Gi + veh rats and control rats (**C**). All experimental groups decreased inactive pressing across self-administration, but did not differ in the number of inactive presses made (**D**). Graphs show group means ± SEM. *p<0.05, **p<0.01.

During self-administration, there was a main effect of training day (F_(13,195)_=7.74, p<0.001), treatment (F_(2,15)_=9.38, p=0.002), and a treatment x training day interaction (F_(26,195)_=2.18, p=0.001) on infusions earned. CNO administration in Gi-DREADD rats decreased the number of infusions earned across training relative to Gi-DREADD rats treated with vehicle (p=0.001) and controls (p=0.003, Figure 1B). Analysis of the interaction indicated that Gi-DREADD rats that received CNO earned fewer infusions than the control group on days 8 and 11-14, whereas they earned fewer infusions than the Gi-DREADD vehicle group on days 2, 4, 6-7, and 9-11 (Figure 1B). Similar results were observed for active lever presses, while no treatment effects were found for inactive lever presses (see Supplement).

### Chemogenetic inhibition alters cue-induced reinstatement of cocaine-seeking

Following self-administration and instrumental extinction, rats experienced repeated cue-induced reinstatement sessions after injection (i.p.) of vehicle, 1mg/kg CNO, and 3mg/kg CNO (order counterbalanced). After each session, rats underwent ∼2 days of re-extinction before the next test day (Figure 2A). An ANOVA with within-subjects factors of test day (extinction vs reinstatement) and CNO dose (vehicle vs 1 mg/kg vs 3 mg/kg) and between subject factors of treatment group during training (control + CNO vs Gi + veh vs Gi + CNO) and sex (male=9 vs female=9) was performed. There was no overall effect of sex on active (F_(1,12)_=2.01, p=0.18) or inactive presses (F_(1,12)_=0.38, p=0.848) made during reinstatement, so data were collapsed for further analysis.

**Figure 2.**
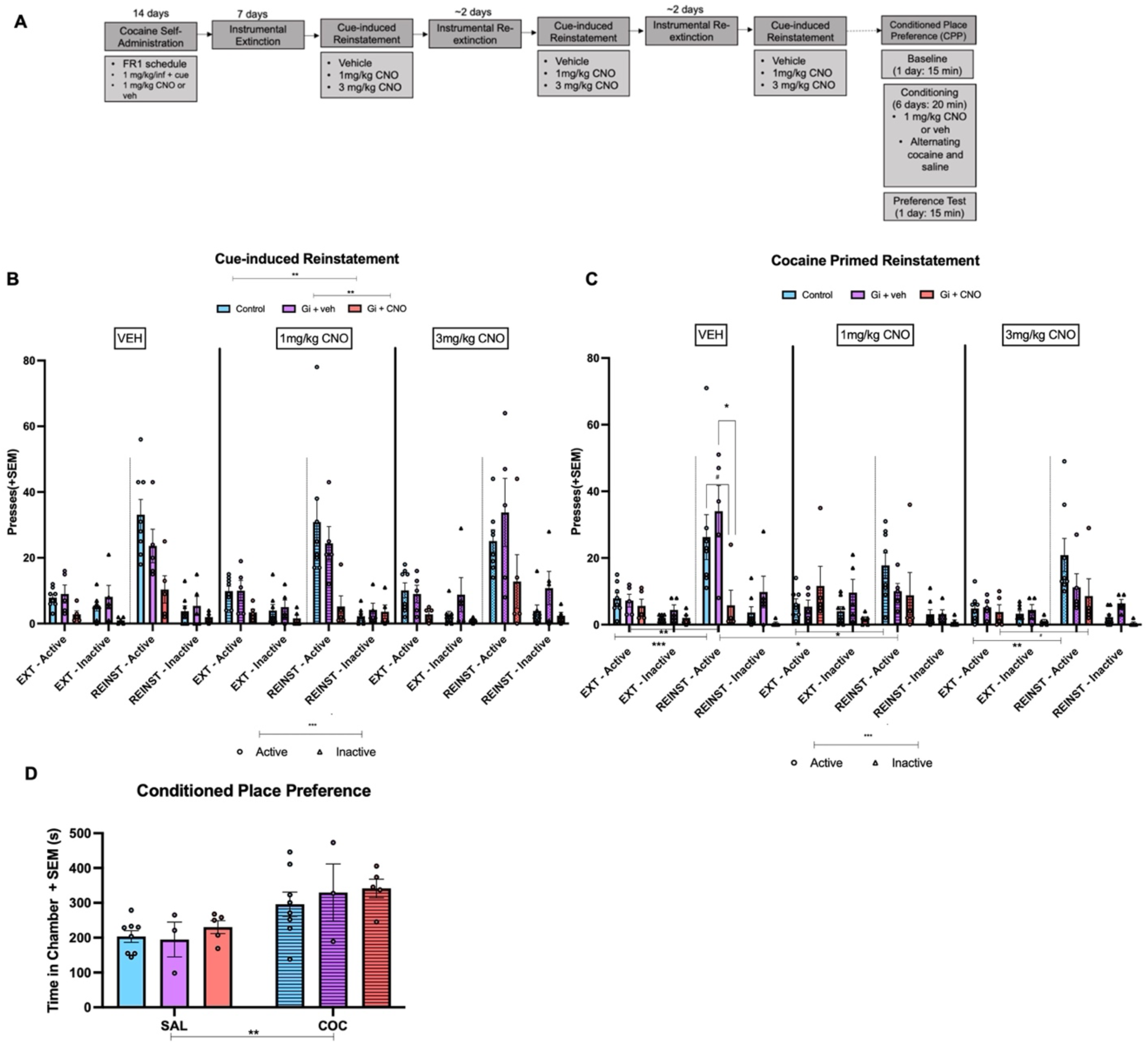
Chemogenetic manipulation affects ability of cues to induce relapse-like behavior, but not cocaine’s ability to induce seeking or a place preference. Rats (n=18, Control=8, Gi + veh=5, Gi + CNO=5) underwent extinction followed by both cue-induced and cocaine primed reinstatement sessions in which they were administered injections of vehicle, 1 mg/kg CNO, or 3 mg/kg CNO (order counterbalanced). A subset of rats (n=16, Control=8, Gi + veh=3, Gi + CNO=5) underwent CPP where they received systemic vehicle or 1 mg/kg CNO during conditioning. Experimental timeline **(A)**. During cue-induced reinstatement, Gi + CNO rats made fewer presses than Gi + veh and control rats, but there was no effect of in-session CNO treatment for any group. Control and Gi +veh rats showed reinstatement to the formerly cocaine-paired cue, while Gi+CNO rats did not **(B)**. During cocaine-primed reinstatement, control rats made more active presses during reinstatement than during extinction during all sessions, Gi + veh rats made more reinstatement presses than extinction presses following 1 mg/kg CNO, and Gi + CNO trended to make more pressing during reinstatement following the 3 mg/kg CNO dose. Following in-session vehicle, Gi + CNO rats made fewer presses than Gi + veh and control rats, but 1 mg/kg CNO treatment in the Gi + veh group blunted cocaine primed reinstatement relative to in-session vehicle. **(C)** On CPP test day, control rats, Gi + veh rats, and Gi + CNO rats spent more time in the cocaine-paired chamber compared to the saline chamber **(D)**. Graphs show group means ± SEM. #p<0.1, *p<0.05, **p<0.01, ***p<0.001.

Analysis of active pressing yielded a main effect of test day (F_(1,15)_=79.24, p<0.001) where generally, rats made more active presses during the reinstatement session than during the day of instrumental extinction that preceded the session (Figure 2B). There was also a main effect of treatment group (F_(2,15)_=7.04, p=0.007) on active pressing, but no effect of in-session CNO dose (F_(2,30)_=0.037, p=0.96). Across both extinction and reinstatement sessions, previous administration of CNO in Gi-DREADD rats attenuated pressing compared to Gi-DREADD rats that received vehicle during training (p=0.009) and control rats (p=0.003, Figure 2B).

There was a test day x treatment group interaction (F_(2,15)_=9.88, p=0.002) and post hoc analysis revealed that both control animals and Gi-DREADD animals that received vehicle during training made more active presses during reinstatement than extinction (p<0.001, p<0.001; respectively), while Gi-DREADD animals that received CNO during training did not (p=0.47, Figure 2B). Thus, rats that experienced inhibition of the VTA DA projection to LA during self-administration showed an impaired ability to reinstate cocaine-seeking in response to the formerly cocaine-paired cue. During all reinstatement sessions, animals discriminated between the active and inactive lever (F_(1,15)_=86.27, p<0.001), but there were no effects of test day (F_(1,15)_=0.032, p=0.86), treatment group during training (F_(2,15)_=1.89, p=0.18), or CNO dose (F_(2,30)_=1.20, p=0.31) on inactive pressing, which remained low across all extinction and reinstatement days.

Cocaine primed reinstatement examines the ability of the cocaine US to elicit drug-seeking, independently of the CS. There was no overall effect of sex on active (F_(1,12)_=0.054, p=0.82) or inactive (F_(1,12)_=1.43, p=0.25) presses made during cocaine primed reinstatement so data were combined. ANOVAs were performed as described above and a main effect of test day (F_(1,15)_=26.11, p<0.001) and a main effect of in-session CNO dose (F_(2,30)_=3.75, p=0.035) on active presses were found, while treatment during training had no effect on behavior (F_(2,15)_=1.31, p=0.299). There was a treatment during training x test day interaction (F_(2,15)_=5.74, p=0.014) as well as a CNO dose x test day interaction (F_(2,30)_=2.93, p=0.010) and a three-way interaction (F_(4,30_)=3.93, p=0.042).

Post hoc analyses of these interactions revealed an interesting pattern. For the control group, the effect of test day was present at all CNO doses as pressing during reinstatement was greater relative to pressing during extinction (vehicle: p<0.001, 1mg/kg: p=0.028, 3 mg/kg CNO: p=0.003). Gi-DREADD rats that received vehicle during training pressed more during reinstatement than extinction following vehicle treatment (p=0.0022). Gi-DREADD rats that received CNO during training trended to make more active presses during reinstatement than during extinction following 3 mg/kg CNO (p=0.092, Figure 2C). Treatment group differences exclusively emerged during reinstatement following in-session vehicle where the Gi-DREADD animals that received CNO during training made significantly fewer active presses than Gi-DREADD animals that received vehicle during training (p=0.042) and trended to make fewer presses than controls (p=0.069, Figure 2C). Gi-DREADD animals that received vehicle during training did not maintain this level of active presses as 1mg/kg CNO during reinstatement blunted active lever responding relative to in-session vehicle (p=0.041). During reinstatement, animals made more active presses than inactive presses (F_(1,15)_=86.26, p<0.001), but inactive pressing behavior during reinstatement was not affected by test day (F_(1,15)_=0.032, p=0.86), CNO dose (F_(2,30)_=1.20, p=0.31), or treatment during training (F_(1,15)_=1.89, p=0.18). Together, the results suggests that inhibition of VTA DA input during cocaine primed reinstatement may decrease seeking behavior in Gi-DREADD animals that did not experience inhibition during training.

An important confirmation of our DREADD strategy was to ensure that systemic CNO administration reduced neuronal activity in the LA of Gi-DREADD expressing TH-Cre animals. We examined c-Fos immunoreactivity as an indicator of neuronal activity after a 1 mg/kg CNO injection followed by a 10 mg/kg cocaine prime (see Supplement). The number of c-Fos+ cells in the LA was lower in the Gi-DREADD group that received CNO compared to those expressing the control virus that received CNO (t(6)=2.47, p=0.049), indicating that CNO in Gi-DREADD rats reduced the number of active cells following cocaine exposure (Supplemental Figure 8).

### Chemogenetic inhibition does not affect cocaine’s rewarding effects

Inhibition of the VTA DA to LA pathway during training appears to alter cocaine cue associative learning to weaken self-administration acquisition and dampen cue-induced reinstatement. These experiments do not rule out that inhibition may impact the rewarding effects of cocaine. To examine whether inhibiting DA inputs to the LA altered cocaine reward/contextual learning, we used the same chemogenetic strategy as above during a conditioned place preference paradigm (CPP). This assay allows us to distinguish between spatial learning and the cue-dependent processes underlying acquisition of self-administration. Comparing time spent in chamber (saline/unpaired vs cocaine/paired) between treatment groups during conditioning (control vs Gi + veh vs Gi + CNO) revealed no main effect of treatment (F_(2,26)_=0.65, p=0.53). After conditioning, control rats, Gi-DREADD rats that received vehicle, and Gi-DREADD rats that received CNO spent more time in the cocaine paired chamber than the saline chamber (F_(1,26)_=14.03, p=0.009, Figure 2F), suggesting that inhibiting VTA DA input to the LA does not impact rewarding properties of cocaine or the ability to learn a contextual association.

### Cocaine self-administration and cue-induced reinstatement are not affected by inhibition of noradrenergic input

Tyrosine hydroxylase (TH) is the rate-limiting enzyme in the synthesis of catecholamines including dopamine and norepinephrine^20^. Our approach targets dopaminergic VTA neurons that project to the LA, but does not rule out the contribution of other TH+ cell populations such as noradrenergic input from the locus coeruleus^21,22^. To determine if this population contributed to the inhibition effects on behavior, we infused a retrograde virus expressing the hM4Di DREADD under the synthetic dopamine beta hydroxylase PRSx8 promoter into the LA of male and female rats. Rats received 1mg/kg CNO injections (i.p.) prior to each cocaine self-administration session for 10 days. We compared the effect of CNO in the PRSx8 group to the inhibition and vehicle groups from the previous Gi-DREADD experiments in TH-Cre rats. There was a main effect of treatment during training (F_(2,17)_=6.20, p=0.0095), training day (F_(2.42, 41.17)_=8.76, p=0.0003), and a treatment x training day interaction (F_(18,153)_=2.85, p=0.0003) on the number of infusions earned (Figure 3B). Similarly, there was an effect of treatment (F_(2,17)_=3.66, p=0.048), training day (F_(3.06, 52.03)_=4.23, p=0.0091), and a treatment x training day interaction (F_(18, 153)_=2.09, p=0.0087) on active presses (Figure 3C). There were no effects of treatment (F_(2,17)_=0.45, p=0.64) or training day (F_(1.60, 27.20)_=3.08, p=0.072) on inactive presses, though inactive pressing trended to decrease across training (Figure 3D). Despite having noradrenergic input silenced, PRSx8 animals performed similarly to the TH-Cre vehicle-treated group and earned more infusions (p=0.008) and made more active presses (p=0.10) than the TH-Cre inhibition animals.

**Figure 3.**
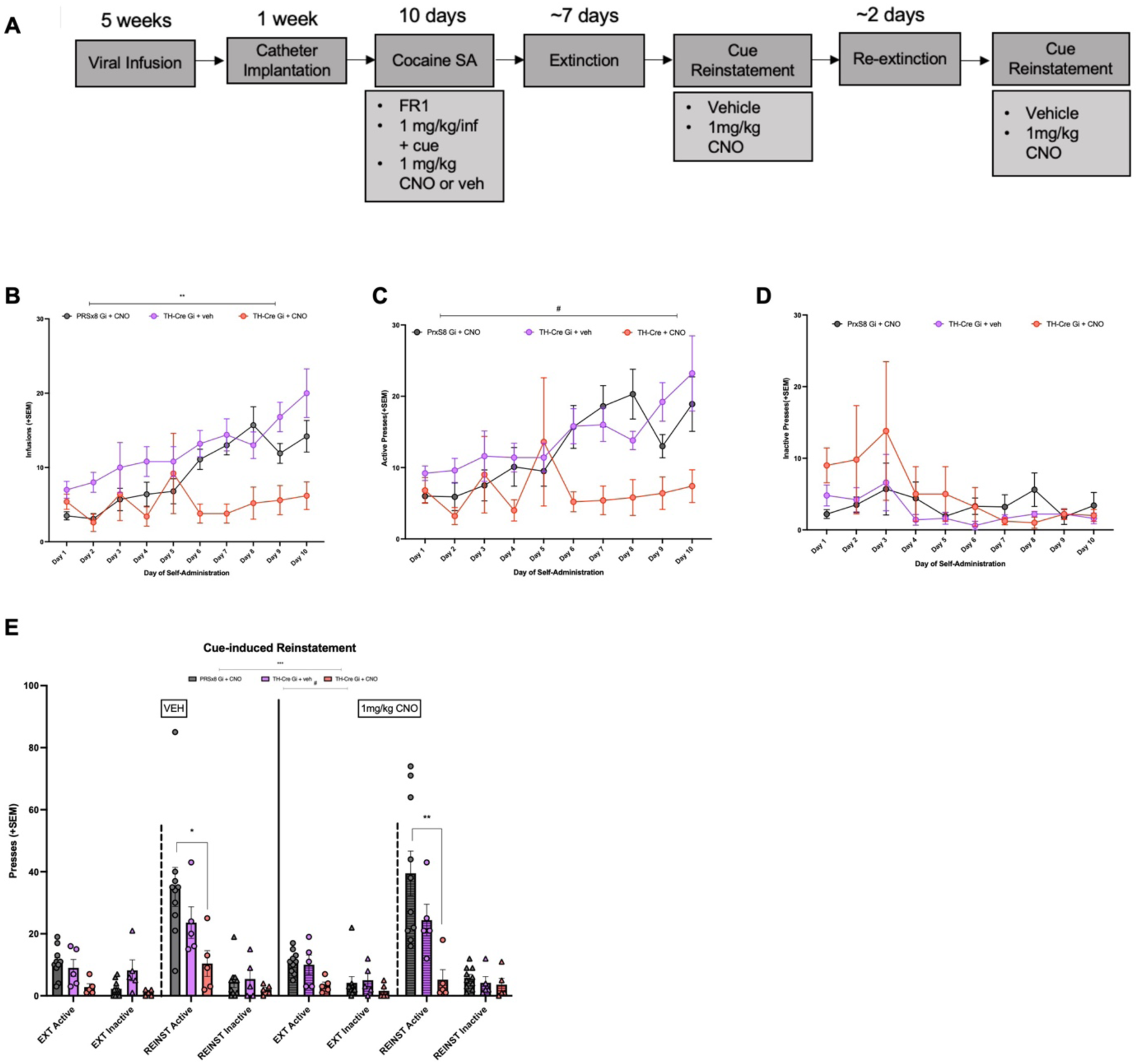
Inhibition of norepinephrine input to LA does not affect behavior. Rats (n=10) were bilaterally infused in the LA with a retrograde AAV expressing the h4MDi inhibitory DREADD receptor under the PRSx8 promoter to infect noradrenergic cells projecting to the region. Experimental timeline **(A)**. PRSx8 Gi-DREADD animals were treated with 1 mg/kg CNO (i.p.) to inhibit norepinephrine input during cocaine self-administration. Their behavior was compared to TH-Cre rats expressing the Gi-DREADD treated with either vehicle (n=5) or CNO (n=5). PRSx8 Gi + CNO rats earned more cocaine infusions than TH-Cre Gi + CNO rats **(B)**. PRSx8 Gi + CNO rats tended to make more active lever presses that TH-Cre Gi + CNO rats **(C)**. There were no group differences in inactive lever presses made during cocaine self-administration **(D)**. Following vehicle and CNO treatment during cue-induced reinstatement, PRSx8 Gi + CNO animals made more active presses than the TH-Cre Gi + CNO rats **(E)** Graphs show group means ± SEM. #p<0.1, *p<0.05, **p<0.01, ***p<0.001.

Following extinction, rats had counterbalanced sessions of cue-induced reinstatement where they received injections of vehicle and 1mg/kg CNO. There was a main effect of test day (F_(1,17)_=24.67, p<0.001) such that active pressing was higher during reinstatement than extinction (Figure 3E). There was no effect of in-session CNO dose (F_(1,17)_=0.018, p-0.89), but there was an effect of treatment during training (F_(2,17)_=10.25, p=0.0012) and a treatment x test day interaction (F_(2,17)_=4.93, p=0.021, Figure 3E). Following vehicle and CNO, PRSx8 animals made more presses than TH-Cre animals that had received CNO during training (p=0.029, p=0.001, respectively). There was no effect of test day (F_(1,17)_=0.43, p=0.52), treatment (F_(2,17)_=1.37, p=0.28) or in-session dose (F_(1,17)_=0.037, p=0.85) on inactive presses. Again, behavior of the PRSx8 group was similar to that of vehicle-treated TH-Cre animals.

### Projection-specific chemogenetic inhibition weakens acquisition of cocaine self-administration

Delivery of the Cre-dependent retrograde virus containing hM4Di receptors results in expression in TH+ VTA cells projecting to the LA, but this chemogenetic protein may be taken up by other TH+ populations innervating the LA located in substantia nigra, periaqueductal gray, or dorsal raphe^23^. To probe VTA to LA input, we employed a strategy where virus was infused into the VTA and CNO was delivered to terminals in the LA. Viruses expressing hM4Di (inhibitory) receptors, hM3Dq (excitatory) receptors, or an EGFP control were infused into the VTA and infusion cannula were implanted over the LA. CNO or ACSF vehicle was infused via cannula prior to the start of each session to examine the effects of inhibition and excitation of the VTA to LA projection on acquisition of cocaine self-administration. 48hr before cocaine self-administration, animals had an initial 6hr food training session to learn the instrumental response to earn reinforcers. There were no group differences in the number of sucrose pellet reinforcers earned (F_(3,77)_=0.29, p=0.83), active presses (F_(3,77)_=0.32, p=0.81), or inactive presses (F_(3,77)_=1.13, p=0.34, Supplemental Figure 6).

To avoid tissue damage from repeated microinfusions, self-administration was limited to 5 days. Males (n=39) and females (n=33) were tested, but there was no effect on sex on infusions earned (F_(1,60)_=0.70, p=0.41), active presses (F_(1,60)_=0.54, p=0.47), or inactive presses (F_(1,60)_=0.006, p=0.94) so data were combined. We also determined if there were group differences within our controls. Post hoc analysis showed that control-expressing animals that received vehicle (n=8) did not differ from control-expressing animals that received CNO (n=11) in terms of infusions, active presses, or inactive presses so they were combined into one control group. Rats expressing the Gi-DREADD virus that received vehicle (n=10) had similar numbers of infusions, active presses, and inactive presses as rats expressing the Gq-DREADD virus that were given vehicle (n=17) so they were combined into one DREADD control group. During self-administration, there was a main effect of treatment (F_(3,68)_=4.79, p=0.004) and a main effect of training day (F_(4,272)_=20.99, p<0.001) on the number of cocaine infusions earned. Gi-DREADD rats that received CNO earned fewer infusions than the Gq-DREADD rats that received CNO (p=0.003). Gi-DREADD rats given CNO showed a trend to earn fewer infusions than the control group (p=0.063) and the DREADD group (p=0.10, Figure 4B). Analysis of active presses revealed main effects of treatment (F_(3, 68)_=4.31, p=0.008) and training day (F_(4,272)_=9.09, p<0.001). Post hoc tests of treatment found that the Gi-DREADD rats that received CNO made fewer presses than the Gq-DREADD rats that received CNO (p=0.008) and the controls (p=0.046). Gi-DREADD rats that received CNO were trending to make fewer active presses than DREADD group rats (p=0.08, Figure 4C). There was no effect of treatment (F_(3,68)_=1.33, p=0.27) or training day (F_(4,272)_=0.56, p=0.64) on inactive pressing (Figure 4D).

**Figure 4.**
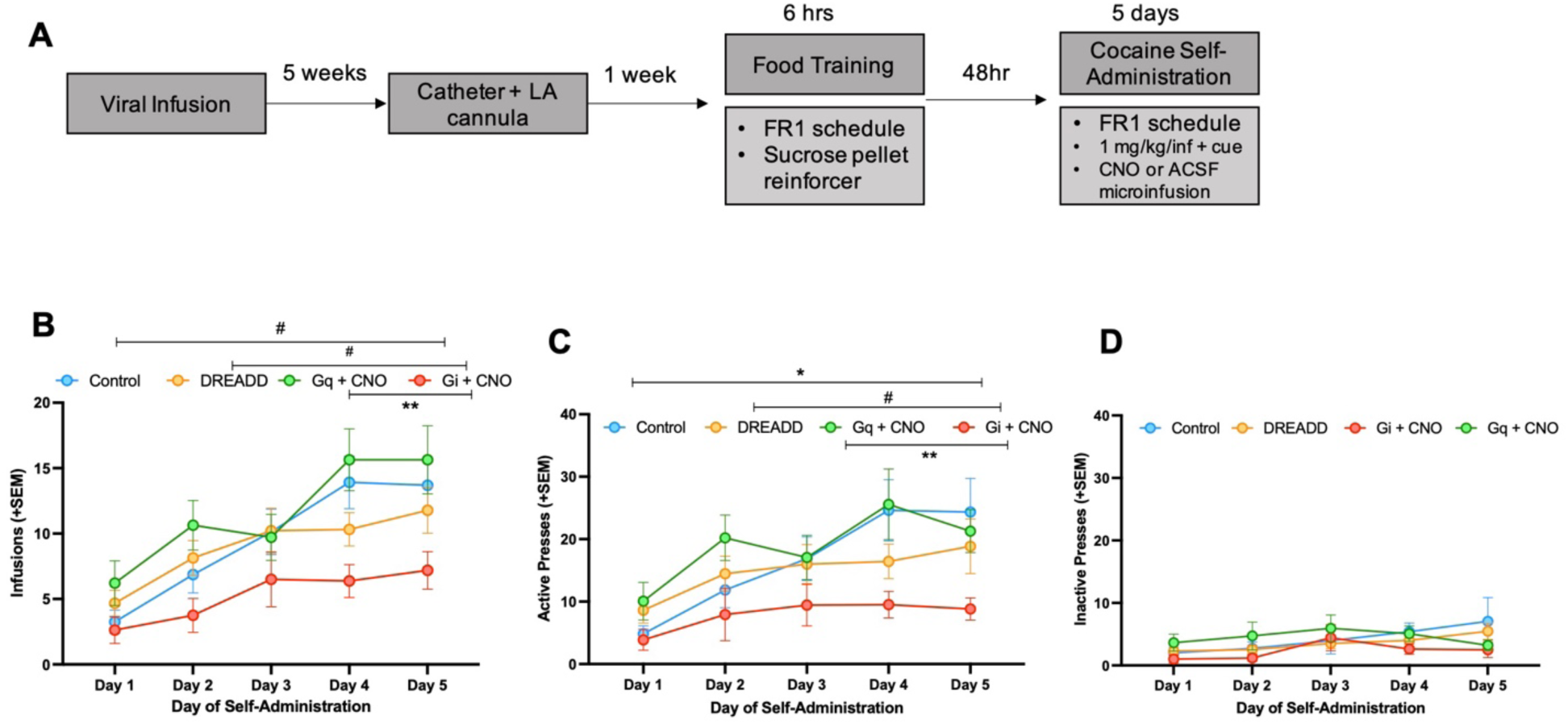
Inhibition of VTA to LA projection weakens acquisition of cocaine self-administration. Rats (n=72, Control=19, DREADD=27, Gi + CNO=15, Gq + CNO=11) were trained to self-administer cocaine on a FR1 schedule and received LA microinfusions of either CNO or ACSF vehicle 5 min prior to each session. Experimental timeline **(A)**. Gi + CNO rats earned fewer cocaine infusions than Gq + CNO rats and trended to earn fewer infusion than DREADD and control rats **(B)**. Gi + CNO rats made fewer active presses relative to Gq + CNO and control rats and trended to make fewer active presses than DREADD rats **(C)**. Experimental groups did not differ in the number of inactive presses made across training **(D)**. Graphs shows group means ± SEM. #p<0.01, *p<0.05, **p<0.01.

### Prior chemogenetic excitation strengthens cue-induced reinstatement, but manipulation does not affect cocaine primed reinstatement

To determine if manipulation of the VTA to LA pathway during self-administration affected relapse-like behavior, animals were tested on cue-induced reinstatement. Males (n=39) and females (n=33) were tested, but there was no effect of sex on active (F_(1,67)_=1.18, p=0.28) or inactive presses (F_(1,67)_=2.98, p=0.089) made during reinstatement. Control animals that received vehicle and control animals that received CNO during training were combined into one group as post hoc analysis revealed no significant difference between the groups. Both Gi-DREADD and Gq-DREADD groups that received vehicle during training were collapsed into a one group as post hoc analysis showed no differences in their behavior.

To assess active lever pressing during cue-induced reinstatement, we performed an ANOVA with within subjects factor of test day (extinction vs reinstatement) and between subjects factor of treatment group during training (control vs DREADD vs Gi + CNO vs Gq + CNO). Analysis yielded a main effect of test day (F_(1,75)_=76.42, p<0.001), a main effect of treatment group (F_(3,75)_=3,19, p=0.028), and a test day x treatment group interaction (F_(3,75)_=2.82, p=0.045). All treatment groups made more active presses during reinstatement than during extinction, suggesting that exposure to the previously drug-paired cue triggered cocaine-seeking in all animals. While there were no group differences during instrumental extinction, differences emerged during cue-induced reinstatement. Gq-DREADD animals that received CNO during training made more active presses than animals in the DREADD group (p<0.001) and Gi-DREADD animals that received CNO during training (p=0.0045, Figure 5B). During reinstatement, more active presses were made than inactive presses (F_(3,204)_=69.85, p<0.001). but there were no differences between treatment groups for number of inactive presses made (F_(3,75)_=0.55, p=0.65).

**Figure 5.**
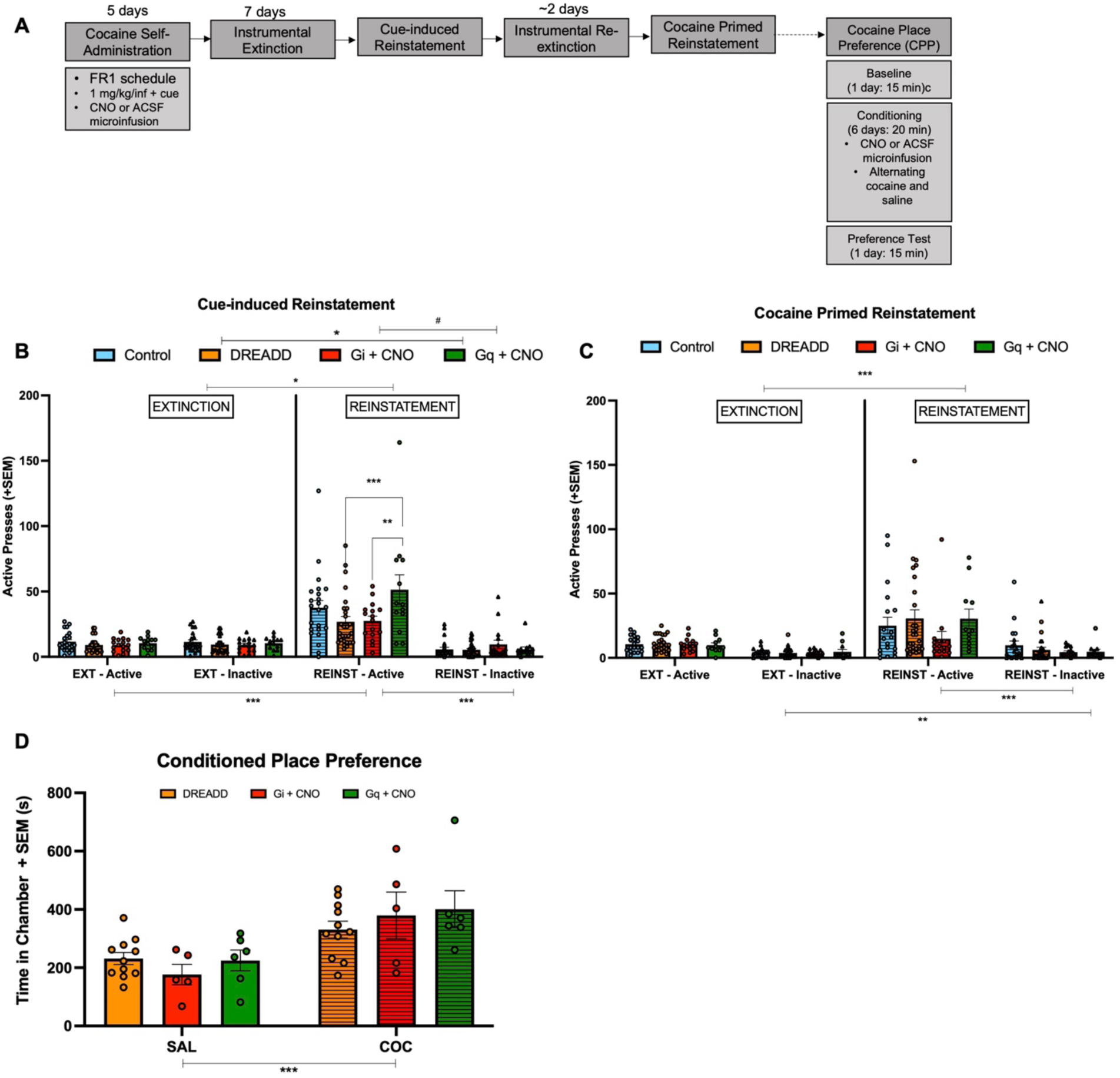
Prior excitation of VTA to LA pathway strengthens cue-induced reinstatement, but manipulation does not affect cocaine-primed reinstatement or cocaine place preference. Rats (n=72, Control=19, DREADD=27, Gi + CNO = 15, Gq + CNO = 11) underwent cue-induced and cocaine primed reinstatement. Rats (n=23, DREADD=11, Gi + CNO=6 Gq + CNO=6) underwent CPP involving LA microinfusions of CNO or vehicle during conditioning. Experimental timeline **(A)**. Gq + CNO rats made more active presses than DREADD group rats and Gi + CNO rats **(B)**. Treatment during self-administration did not impact reinstatement to a cocaine prime **(C)**. After conditioning, DREADD, Gi + CNO, and Gq + CNO rats spent more time in the cocaine-paired chamber than the saline-paired chamber **(D)**. Graphs show group means ± SEM. #p<0.01, *p<0.05, **p<0.01, ***p<0.001.

It was important to confirm that CNO affected neuronal activity in the LA of DREADD expressing rats. We examined c-Fos immunoreactivity following a cue-induced reinstatement session since c-Fos expression serves as an indicator of activity (see Supplement). CNO or vehicle microinfusions were administered into the LA of Gi-DREADD-expressing, Gq-DREADD-expressing and control rats prior to the session. Analysis of c-Fos+ cells yielded a main effect of treatment (F_(2,13)_=5.78, p=0.016). Gi-DREADD rats that received a CNO microinfusion had fewer c-Fos+ cells than DREADD controls (p=0.039) and Gq-DREADD animals that received CNO (p=0.022). CNO administration in the inhibition group reduced the number of active LA cells following cue-induced reinstatement (Supplemental Figure 9).

We examined if manipulation during training impacted cocaine-primed reinstatement. Both sexes were used, but there was no effect of sex on the number of active (F_(1,60)_=1.75, p=0.19) or inactive presses (F_(1,60)_=2.38, p=0.13). Post hoc analysis showed no group difference between control animals that received vehicle or those that received CNO so they were pooled into one control group. Post hoc comparisons revealed no difference between Gi-DREADD animals that received vehicle and Gq-DREADD animals that received vehicle so animals were combined into one DREADD group.

There was a main effect of test day (F_(1,68)_=13.39, p<0.001), but no effect of treatment during training on active presses during cocaine primed reinstatement (F(3,68)=1.38, p=0.095), suggesting that all animals behaved similarly during cocaine primed reinstatement whether or not they experienced inhibition or excitation during training. However, in general, responding to the cocaine prime was low. During cocaine primed reinstatement, rats made more active presses than inactive presses (F_(1,68)_=23.28, p<0.001). Examining inactive presses on their own revealed a main effect of test day on inactive presses (F_(1,68)_=75.24, p<0.001) such that inactive pressing was greater during reinstatement than extinction. However, there was no effect of treatment during training (F_(3,68)_=0.038, p=0.99) indicating that all rats made a similar number of inactive presses regardless of prior VTA DA to LA projection manipulation.

### Chemogenetic manipulation of the VTA to LA projection does not impact cocaine’s rewarding effects

Our results suggest chemogenetic manipulation of the VTA to LA projection affects cue learning, but we again wanted to determine potential effects on cocaine reward/contextual learning using CPP. An ANOVA with within subjects factor of time spent in chamber (saline (unpaired) vs cocaine (paired)) and between subjects factor of treatment during conditioning (control vs DREADD vs Gi + veh vs Gi + CNO) was performed. There was no main effect of treatment during conditioning on time spent in each chamber (F_(2,38)_=0.38, p=0.69). Following conditioning, all treatment groups spent more time in the cocaine paired chamber than the saline paired chamber (F_(1,38)_=20.88, p<0.001, Figure 5D).

## Discussion

Here, we examined the effects of chemogenetic manipulation of the VTA DA to LA projection during cocaine self-administration on acquisition and reinstatement. Acquisition of cocaine self-administration was disrupted when animals had DA input from the VTA to LA inhibited. When this projection is silenced during self-administration, animals later show a reduction in cocaine-seeking elicited by the previously cocaine-paired cue. Conversely, excitation of the projection boosts subsequent cue-induced reinstatement. Inhibiting this projection did not impact the ability of cocaine to elicit a place preference or to induce reinstatement in response to a priming injection. Thus, these results suggest that the VTA DA to LA projection plays an important role in cocaine cue associative learning/reinforcement of instrumental responding that is critical for acquisition of cocaine self-administration, but does not alter rewarding effects of cocaine or spatial reward learning.

Previous research demonstrated that cues facilitate acquisition of self-administration and that these cues do not change drug-reinforcing effects, but rather modulate drug-seeking behavior^24–26^. Our findings are consistent with these studies as we showed that rats undergoing inhibition of the VTA to LA pathway still self-administered cocaine, but at much lower levels than controls. The LA is critical for cocaine-cue associative learning and interfering with DA input to this region disrupted the ability to associate the lever response with the discrete cue and effects of cocaine, which weakened the ability to acquire cocaine self-administration.

Assessing cue-induced reinstatement helped determine how inhibition during self-administration affected the expression of cue learning. Cue-induced reinstatement is a commonly used model of relapse-like behavior that probes the strength of cue associations^27–29^. Lesioning the basolateral amygdala (BLA), which contains the LA, has no effect on responding during cocaine self-administration, but abolishes cue-induced reinstatement^27,30^, indicating the region has a specific role in mediating cue-dependent behaviors. There was similar specificity for rats that experienced excitation of the VTA to LA projection during 5 days of training. These animals did not have increased responding during self-administration, but had the highest levels of cue-induced reinstatement, implying that excitation strengthened drug-cue associative learning.

When rats had DA projections to the LA inhibited for 14 days, we also found a reduction in active pressing in response to the formerly drug-paired cue, further highlighting the projection’s role in cue-dependent behavior. However, low levels of self-administration in this group likely limits the amount of cue-induced reinstatement that could be observed.

Comparing cocaine primed and cue-induced reinstatement can disentangle the effects of cocaine itself from the cue’s ability to trigger drug-seeking. In general, manipulation of the VTA DA projection to the LA during self-administration did not affect later cocaine primed reinstatement, although there was a low level of responding. The amount of cocaine intake during self-administration and the magnitude of subsequent cocaine-induced reinstatement are related^31,32^. Rats that have more access to cocaine via longer self-administration sessions or higher doses of cocaine are more susceptible to cocaine primed reinstatement^33,34^. This pattern may explain why reinstatement to the cocaine prime was not robust in our experiments.

We observed a significant difference between TH-Cre animals expressing the Gi-DREADD that received vehicle versus CNO during training. This likely reflected the difference seen during self-administration where the Gi-DREADD animals treated with vehicle earned more infusions than the Gi-DREADD animals treated with CNO. Within the group of TH-Cre animals expressing the Gi-DREADD that received vehicle during training, in-session CNO treatment tended to blunt responding to a cocaine prime. Others have shown that chemogenetic inhibition of all VTA DA neurons attenuates cocaine primed reinstatement^18^. Thus, in the absence of inhibition-related plasticity changes during training, acutely altering the VTA to LA projection may contribute to weakening cocaine primed reinstatement.

As manipulating the VTA DA to LA projection during self-administration impacted acquisition and cue-induced reinstatement, but not cocaine primed reinstatement, it seems to be a pathway heavily involved in cocaine cue associative learning. Additionally, neither inhibition nor excitation of the VTA to LA projection during conditioned place preference (CPP) affected the animals’ contextual learning or ability to develop a preference for the cocaine paired chamber. Amygdala lesions block cocaine-induced CPP^35^, but silencing just the VTA DA projection to the LA left CPP intact, further highlighting its role in associative learning rather than contextual learning or reinforcement.

A limitation of this study is that there are not groups that self-administers saline or undergo cue-free self-administration. A saline self-administration group would help determine if chemogenetic manipulation effects are drug-specific. Groups trained to self-administer cocaine without cues would provide insight into whether the deficit in acquisition was specific to associative learning related to the cue versus the lever pressing action. Although these additional conditions could strengthen our conclusions, replicating our results using two different projection targeting strategies lends support to our findings. Additionally, the dissociable effects of chemogenetic manipulation on cue-dependent behaviors versus spatial reward-related learning provides strong evidence that the VTA DA to LA projection is not contributing to the rewarding effects of cocaine.

Cues paired with repeated drug use drive craving and relapse and here we demonstrated that weakening this input during acquisition of self-administration attenuated relapse-like behavior elicited by cues. Additionally, these experiments provide support for VTA DA as an LA input that encodes the reinforcing effects of cocaine that serve as a US in conditioned cue drug-seeking. Dealing with multiple drug-paired cues and contexts is a frequent obstacle for substance use disorder treatments such as cue exposure therapy, but preclinical work targeting reactivation of US memory to extinguish more than one CS highlights the promise of disrupting US encoding^36^. Our findings present a potential US-relevant pathway in the development of substance use disorder that warrants further investigation.

## Acknowledgements

We would like to acknowledge Brooke Bender, Camryn Forbes, Michael Wright, Kelly Lai, Allison Caswell, and Kacie Barry for assistance with behavioral experiments, surgical procedures, and histology. We would like to thank Jenny Zeak for animal breeding and genotyping and Sierra Stringfield for technical advice. This work was supported by the National Institutes of Health R01DA042029 and R01AA02815.

## Notes

### Competing Interest Statement

The authors have declared no competing interest.

