## Supplemental Material for "The ventral tegmental area dopamine to lateral amygdala projection supports cocaine cue associative learning"

### Methods

#### Viral constructs and delivery

Rats were placed in a stereotaxic frame and given an injection (0.2-0.3 ml) of lidocaine (Henry Schein) to the scalp. For chemogenetic manipulations in TH-Cre rats, a 26-gauge stainless steel injection cannula connected to a Hamilton syringe was used to inject the LA (males in mm from bregma, anterior and posterior (AP): -3.0; medial and lateral (ML): ±5.0; dorsal and ventral (DV): -7.9 mm; females in mm from bregma, AP: -2.8, ML: ±4.8; DV: -7.8 mm) bilaterally (1μl/hemisphere) through a pump (Harvard Apparatus) with retrograde adeno-associated virus (rgAAV) containing a double-floxed, inverted open reading frame (DIO) sequence for the mCherry-tagged hM4Di(Gi-coupled) or mCherry control (hSyn-DIO-hM4D(Gi)-mCherry, hSyn-DIO-mCherry, Addgene). Using the same setup, a custom AAV2retro.PRSx8.HA-hM4D.SV40 (Penn Vector Core) was bilaterally infused into the LA In all other chemogenetic manipulation experiments, the setup was used to inject the VTA (males and females: in mm from bregma AP: -5.5, ML: ± 0.9; DV: -8.20) bilaterally with 1μl/hemisphere adeno-associated virus 2 (AAV2) containing the mCherry-tagged hM4Di (Gi-coupled) or hM3Dq (Gq-coupled) DREADDs or EGFP control (CaMKII**α** -hM4D(Gi)-mCherry, CaMKII**α**-hM3D(Gq)-mCherry, CaMKII**α**-EGFP, Addgene). Injections were at a 0.1 µl/min rate and injection cannula were left in place for 5 min after completion before being withdrawn. More than 5 weeks was allowed between virus injection and training.

### Surgery

#### Catheter implantation

Rats were anesthetized via intramuscular injections of ketamine (90-100mg/kg, Henry Schein) and xylazine (5mg/kg, Butler Schein) and were given subcutaneous injections of the analgesic Rimadyl (5mg/kg, Henry Schein) and 5 mL of Lactated Ringer's solution. Surgical sites were shaved and treated with betadine (povidone iodine, 5%, Henry Schein) and 70% ethanol on all incisions. Rats were implanted with a chronic indwelling intravenous catheter into the right jugular vein as previously described[1].

#### Intracranial cannulation

Following catheterization, rats used in experiments involving intra-LA microinfusions were placed in a stereotaxic frame and received lidocaine (0.2-0.3 ml, Henry Schein) to the scalp. Two 22-gauge guide cannula (cut 11 mm below 8 mm pedestal, PlasticsOne) were implanted bilaterally, aimed 2 mm dorsal to LA (males in mm from bregma, anterior and posterior (AP): -3.0; medial and lateral (ML): ±5.0; dorsal and ventral (DV): -7.9 mm; females in mm from bregma, AP: -2.8, ML: ±4.8; DV: -7.8 mm). Cannula were secured to the skull with screws and OrthoJet dental cement (Lang Dental). Once cured, dummy cannula (C313DC, PlasticsOne) were inserted to prevent obstruction.

#### Post-operative care

For two days following surgery, rats were given Rimadyl (5mg/kg, Henry Schein) subcutaneously. Catheter patency was maintained by daily infusion of 0.1 ml of a 0.9%

sterile saline solution containing Gentamicin (3mg/ml, Henry Schein) and heparin (30 USP/ml; Henry Schein).

**Behavioral Apparatuses**

Experiments were conducted in 24 standard operant conditioning chambers (MedAssociates) using MedPC software (MedAssociates). Each animal underwent all training and testing in the same chamber. Each chamber was equipped with bar floors, a house light, two cue lights above two levers, a tone generator, a head-entry magazine, a sucrose pellet dispenser, and a syringe pump connected to a swiveled leash. All chambers had 2 plexiglass walls with one containing the levers, magazine and cue lights and the opposite wall containing nose-poke apertures. Half of the chambers were equipped with 2 nose poke apertures, while the others were equipped with 5 nose-poke apertures with a removable opaque plexiglass cover. Chambers were contained in sound-attenuating boxes with fans for background noise. Conditioned place preference experiments were conducted in a custom made 60 cm X 30 cm X 30 cm apparatus. The apparatus consisted of 2 25 cm X 30 cm X 30 cm chambers connected by a 10 cm X 30 cm X 30 cm pathway containing 15 cm X 15 cm entryways. During conditioning and testing, chambers differed from one another in odor (vanilla or almond), floor texture (Plexiglass or rough plastic covering), and wall pattern (vertical stripes or horizontal stripes). During conditioning, a 30 cm X 30 cm X 2 cm divider was inserted over the entryway to restrict the animal to a single chamber.

**Histology**

Rats were perfused by first deeply anesthetizing them with sodium pentobarbital (Coventrus, 100 mg/kg i.p.) and then perfusing through the aorta 1X PBS for 5 min followed by 4% paraformaldehyde (Santa Cruz Biotechnology) in 1X PBS, pH 7.4 for 10 min. The brains were extracted, postfixed in 4% paraformaldehyde for 24hr before being transferred to a 30% sucrose solution. Brains were sectioned at 50 μm using a cryostat (Leica). Slices containing the LA or VTA were collected for histology or immunohistochemistry. In the case of histology, slices were mounted on glass slides and coverslipped with Fluoroshield with DAPI mounting media (Sigma-Aldrich). Slides were imaged using an Olympus VS120 slide-scanning microscope to verify virus expression and/or cannula placement. Rats lacking expression of AAV in LA or VTA and those with improperly positioned cannula were removed from the study. For experiments with TH-Cre rats, all groups had equivalent percentages of cells infected in both the LA (percent infected ± SEM: Control: 41.56% ± 9.39; Gi + veh: 39.96% ± 6.22; Gi + CNO: 46.32% ± 4.53) and VTA (percent infected ± SEM: Control: 62.05% ± 4.14.; Gi + veh: 56.58% ± 2.09; Gi + CNO: 58.15% ± 4.30). For the VTA to LA projection manipulation via local microinfusions, all groups had equivalent percentages of cells infected in both the VTA (percent infected ± SEM: Control: 57.54% ± 2.12; Gi + veh: 63.45% ± 1.72; Gq + veh: 63.44% ± 4.19; Gi + CNO: 65.14% ± 3.92; Gq + CNO: 65.983% ± 3.60) and LA (percent infected ± SEM: Control: 54.38% ± 2.23; Gi + veh: 52.39% ± 2.07; Gq + veh: 47.19% ± 2.73; Gi + CNO: 48.39% ± 2.03%; Gq + CNO: 50.89% ± 9.10)

In the PRSx8 promoter Gi-DREADD experiments, animals were sacrificed as described above and sections containing the LA or LC were collected to examine immunoreactivity of the HA-tag associated with the virus. These sections were washed with PBS containing 0.1%

triton X (PBST+) and then incubated in a PBST+ and 5% donkey serum (Millipore Sigma) blocking buffer for 2hr at room temperature. Sections were then placed in anti-HA primary antibody (1:1000 Cell Signaling C29F4) for 48hr at 4ºC. After incubation in primary antibody, sections were washed with PBST+ and moved to wells with secondary antibody in blocking buffer (1:500 Donkey anti-rabbit IgG Alexa Fluor 594, ThermoFisher) for 2hr at room temperature. Sections were washed in PBS, mounted, and coverslipped with Fluoroshield with DAPI mounting medium (Sigma Aldrich). Slides were then scanned using an Olympus VS120 scanning microscope. Within the LA, the mean percent of cells infected was 59.80% ± 2.91, while the mean percent infected was 53.08% ± 3.85 in the LC.

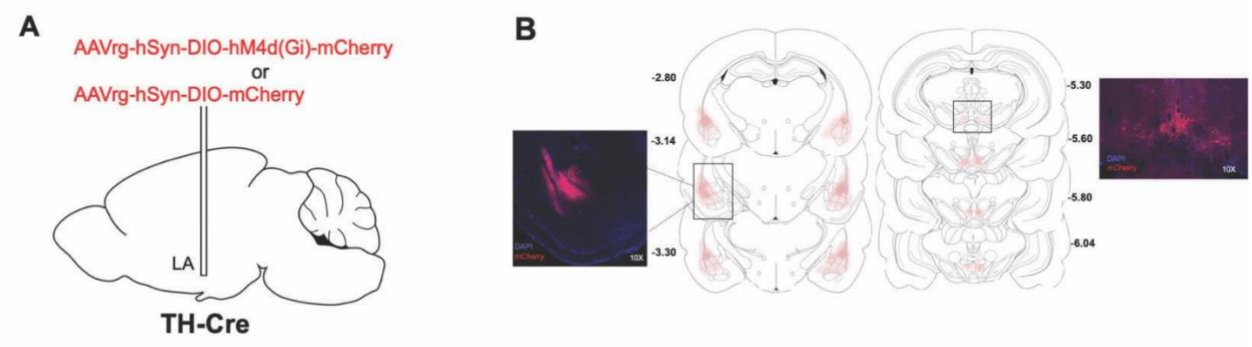


**Supplemental Figure 1. Virus infusion schematic and expression in TH-Cre rats.** Schematic of infusion of retrograde Cre-dependent inhibitory Gi-DREADD virus or mCherry control virus in LA of male and female TH-Cre rats **(A).** Representative images of virus expression (mCherry) and spread in both the LA and VTA **(B)**.

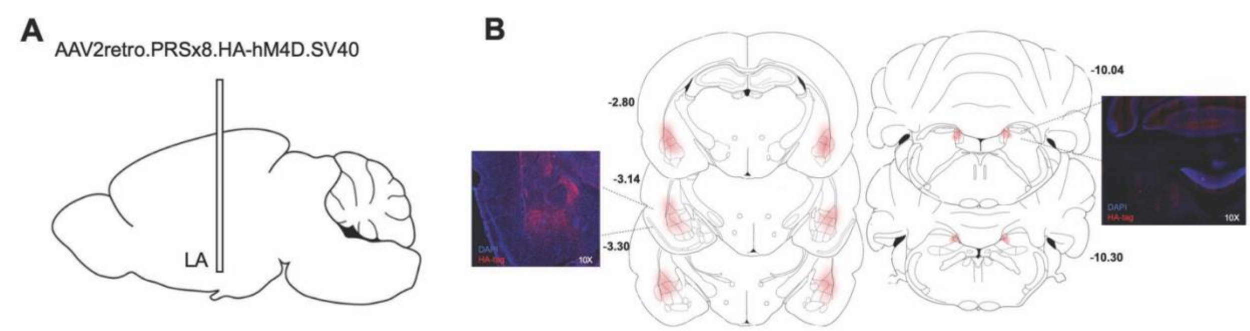


**Supplemental Figure 2**. **Virus infusion schematic and expression.** Schematic of infusion of retrograde PRSx8 promoter Gi-DREADD virus in LA of male and female Sprague Dawley rats **(A)**. Representative images of virus expression (HA-tag) and spread in both the LA and locus coeruleus **(B)**.

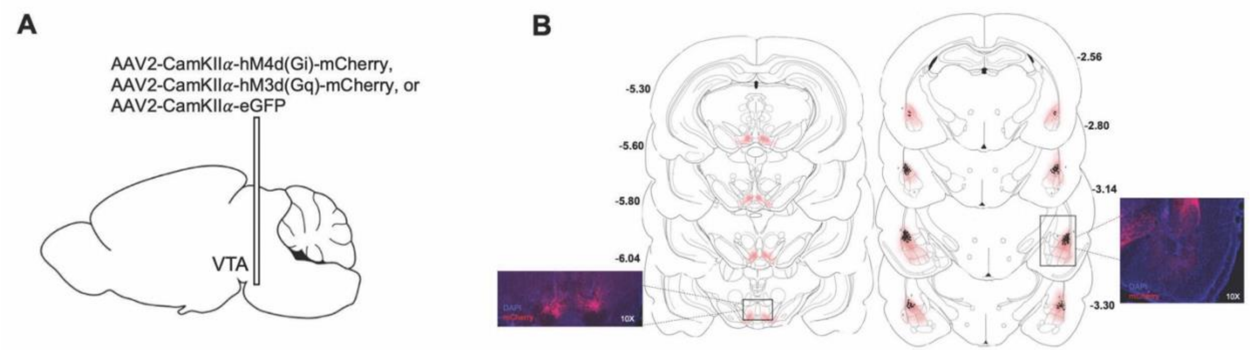


**Supplemental Figure 3. Virus infusion schematic, expression, and cannula placement.** Schematic of infusion of inhibitory Gi-DREADD, excitatory Gq-DREADD, and EGFP control virus in VTA of male and female Sprague Dawley rats **(A)**. Representative images of virus expression (mCherry) and spread in both the VTA and LA. Black circles indicate proper cannula placement, while black “X” mark unsuccessful placements **(B)**.

### c-Fos immunoreactivity

c-Fos immunoreactivity was used as an indicator of activity to determine the efficacy of CNO in DREADD expressing animals. These animals were sacrificed via transcardial perfusion 90 minutes following a final behavioral session. To verify the effects of CNO in TH-Cre rats, animals were sacrificed 90 min following a 10mg/kg cocaine i.p. injection that was preceded by an i.p. injection of CNO. To confirm CNO effects following a local microinfusion, rats received a CNO microinfusion prior to a final cue reinstatement session and were euthanized via cardiac perfusion 90 min after the session. Brains were harvested and sectioned at 50 µm using a cryostat (Leica). Sections were washed with PBS with 0.1% triton (PBST+) and then left in a PBST+ and 5% donkey serum (Millipore Sigma) blocking buffer for 2hr at room temperature. Sections were then incubated in anti c-Fos primary antibody (1:2000, Abcam ABE457) for 48 hrs at 4°C. After primary incubation, sections were again washed with PBST+ and then placed in wells with secondary antibody in blocking buffer (1:500, Donkey anti-rabbit IgG Alexa Fluor 488, Invitrogen Antibodies) for 2hr at room temperature. Sections were then washed in PBS, mounted, and coverslipped with Fluoroshield with DAPI mounting medium (Sigma Aldrich). Slides were scanned using an Olympus VS120 scanning microscope with the DAPI, RFP and GFP channels. c-Fos+ cells were counted with Fiji (ImageJ) software. Three sections per region per animal were counted and averaged to give a cell count per animal

### Results

#### Chemogenetic inhibition during self-administration affects active pressing behavior.

There were main effects of training day ($F_{(13,195)}=4.69$, $p<0.001$) and treatment group ($F_{(2,15)}=7.50$, $p=0.006$), and a training x treatment interaction ($F_{(2,195)}=1.81$, $p=0.013$) on the number of active presses made during cocaine self-administration. Gi-DREADD expressing rats that received CNO made fewer active presses than Gi-DREADD rats that received vehicle ($p=0.002$) and controls ($p=0.010$, see Figure 1C). Post hoc analysis of the interaction revealed Gi-DREADD rats treated with CNO made fewer active presses than Gi-DREADD rats given vehicle on days 2, 4, 6-7, 9, and 11-12. On days 7-9, and 11-12 CNO treatment in Gi-DREADD expressing rats led to fewer active lever presses compared to controls. There was no main effect of treatment condition on inactive press ($F_{(2,15)}$=1.54, p=0.25), but an effect of training day ($F_{(13,195)}$=3,69, p<0.001) as all animals decreased inactive pressing across training (see Figure 1D).

**Prior inhibition of VTA DA input to LA in TH-Cre rats affects early but not late instrumental extinction.**

Self-administration was followed by instrumental extinction to extinguish lever pressing behavior before evaluating the effects of inhibition on the ability of the cocaine-paired cue or cocaine to reinstate drug-seeking behavior. There was a main effect of treatment on active presses made during extinction ($F_{(2,15)}$=4.77, p=0.025) and a main effect of extinction day ($F_{(6,90)}$=12.61, p<0.001, Supplemental Figure 4A). These effects were driven by rats in the control group that made more active presses than Gi-DREADD rats that received CNO (p=0.028) and made more presses than Gi-DREADD rats that were given vehicle (p=0.074). Multiple comparisons showed that the control group made more active presses than the inhibition group on extinction days 1 and 4. However, all groups equally extinguished their pressing behavior by the final extinction day (Supplemental Figure 4A).

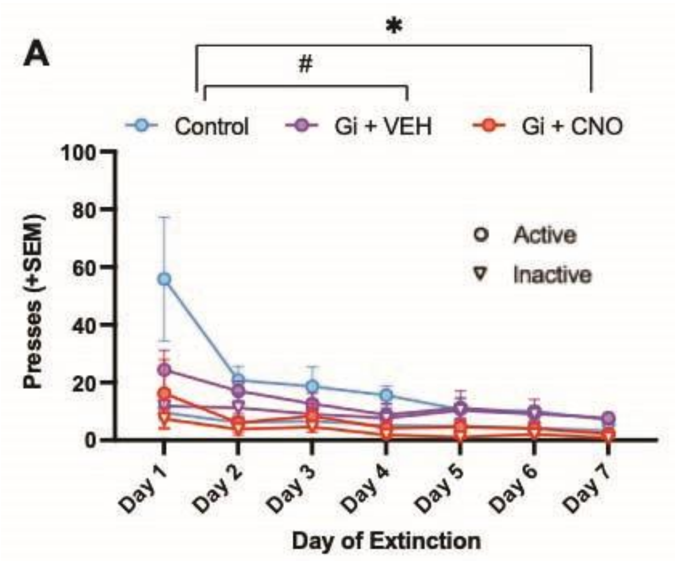


**Supplemental Figure 4. Prior inhibition of VTA DA to LA projection impacts early, but not late instrumental extinction.** Rats (n=16, Control = 8, Gi + veh =3, Gi + CNO =5) underwent instrumental extinction following 14-days of cocaine self-administration. Treatment during self-administration affected the number of active lever presses made during instrumental extinction as control rats made more active presses than Gi-DREADD rats that received CNO and trended to make more active presses than the Gi-DREADD rats that received vehicle. All animals decreased their pressing across training and reached a low level of pressing by the final day **(A)**. Graph shows group means ± SEM. Circles indicate active presses while triangles indicate inactive presses.

### Differences in instrumental extinction behavior found between PRSx8 Gi-DREADD group given CNO and the TH-Cre inhibition group.

Comparing extinction between the PRSx8 Gi-DREADD group, the TH-Cre vehicle treated Gi-DREADD group, and the TH-Cre Gi-DREADD inhibition group revealed a main effect of treatment during training ($F_{(2,21)}=4.34$, $p=0.026$) and a main effect of training day ($F_{(2,65, 55.73)}=10.10$, $p<0.001$**)** on active presses**.** Across extinction, rats in the PRSx8 group made more active presses than rats in the TH-Cre inhibition (Supplemental Figure 5A). Despite this difference, all groups decreased their pressing behavior across extinction and were within extinction criteria by the end of instrumental extinction by making fewer than 15 active presses on average. There were no group differences in the number of inactive presses made during extinction ($F_{(2,21)}$=1.25, p=0.31), but there was an effect of extinction day where all animals decreased inactive presses as extinction progressed ($F_{(3.75, 78.66)}$=4.39, p=0.0036).

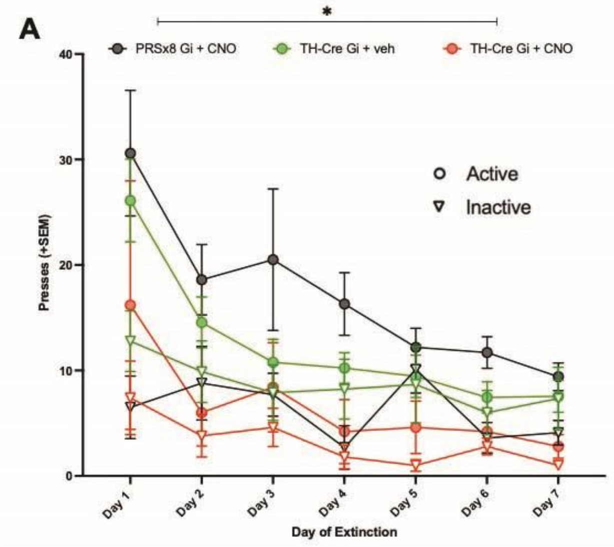


**Supplemental Figure 5. Differences in extinction between PRSx8 Gi-DREADD animals and TH-cre Gi-DREADD inhibition animals.** Rats (n=20, PRSx8 Gi+CNO=10, TH-Cre Gi+veh=5, TH-Cre Gi+CNO=5) underwent 7 days of instrumental extinction following cocaine self-administration. Rats in the PRSx8 Gi+CNO group during training made more active presses across extinction than animals in the TH-Cre+CNO group during training, but all groups (PRSx8 Gi+CNO, TH-Cre Gi+veh, and TH-Cre Gi+CNO) made fewer than 15 presses on average. Inactive pressing was similar for all groups **(A)**. Graphs shows group means ± SEM. *p<0.05. Circles indicate active presses while triangles indicate inactive presses.

### During food training, groups do not differ in number of sucrose reinforcers, active presses, or inactive presses.

48hr before beginning cocaine self-administration paired with pre-session microinfusions of CNO or vehicle, animals had an initial 6hr food training session on an FR1 schedule to learn the appropriate instrumental response to earn reinforcers. No audiovisual cues were presented during the training session. There were no group differences in the number of sucrose pellet reinforcers earned ($F_{(3,77)}$=0.29, p=0.83, Supplemental Figure 6A), active presses ($F_{(3,77)}$=0.32, p=0.81, Supplemental Figure 6B), or inactive presses ($F_{(3,77)}$=1.13, p=0.34, Supplemental Figure 6B).

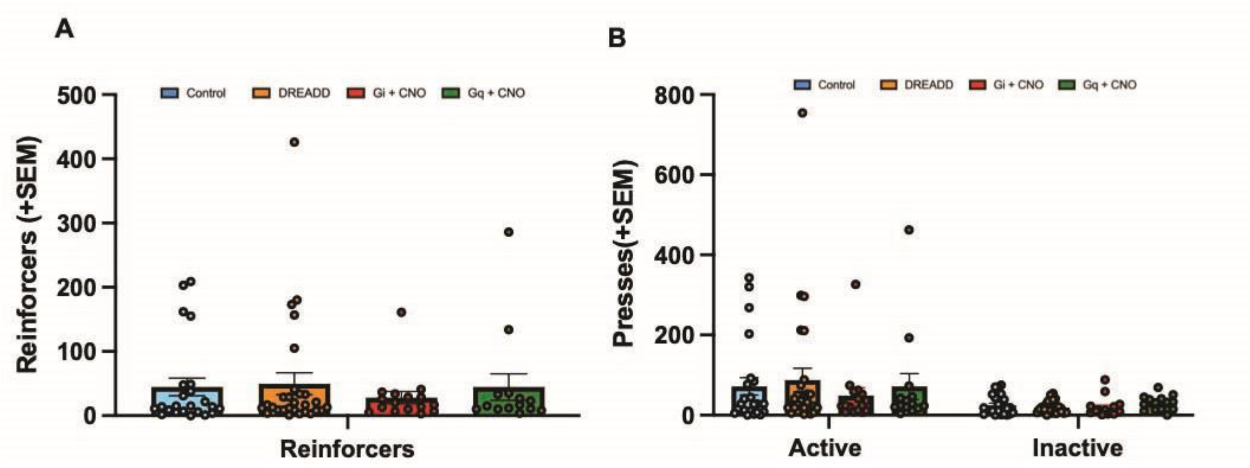


**Supplemental Figure 6. Performance during food training does not differ between groups.** Rats (n=72, Control=19, DREADD=27, Gi + CNO = 15, Gq + CNO = 11) were trained to self-administer sucrose pellets during a 6-hour food training session before any microinfusions occurred. There were no significant group differences in reinforcers **(A)**, active presses **(B)**, or inactive presses **(B)**. Graph**s** shows group means ± SEM.

**Prior chemogenetic inhibition of the VTA to LA pathway does not affect instrumental extinction.**

After cocaine self-administration, animals underwent instrumental extinction to extinguish their active lever pressing behavior before continuing into reinstatement testing. There was no effect of treatment during self-administration ($F_{(3,68)}$=1.13, p=0.34) on the number of active presses made. There was a main effect of extinction day ($F_{(6,408)}$=27.75, p<0.0001) as lever pressing decreased across days of extinction (Supplemental Figure 7A).

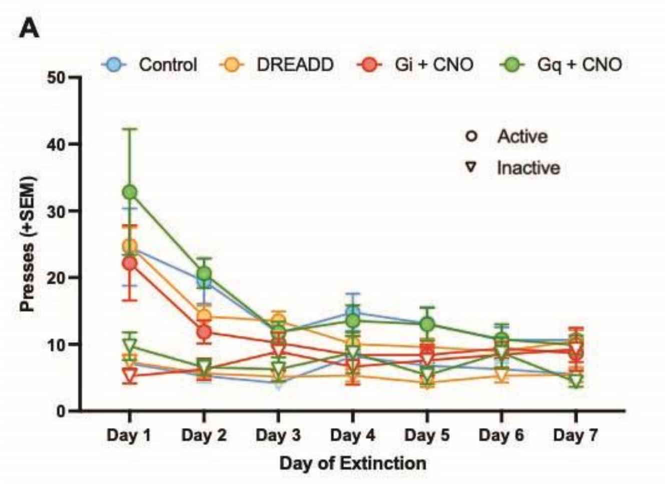


**Supplemental Figure 7. Chemogenetic inhibition of the VTA to LA pathway during self-administration does not impact instrumental extinction.** Rats (n=72, Control=19, DREADD=27, Gi + CNO = 15, Gq + CNO = 11) underwent instrumental extinction following cocaine self-administration. Treatment during cocaine self-administration did not affect pressing behavior during extinction. There was a main effect of extinction day as all rats decreased their lever pressing across extinction days **(A)**. Graph shows group means ± SEM. Circles indicate active presses while triangles indicate inactive presses.

#### Administration of CNO in Gi-DREADD expressing TH-Cre rats impacts c-Fos immunoreactivity following a cocaine prime.

Binding of CNO to the Gi-DREADD receptor triggers hyperpolarization of the cell, inhibition of neurotransmitter release, and neuronal silencing. As a confirmation that systemic CNO administration reduced overall neuronal activity in the LA of Gi-DREADD expressing animals, we examined c-Fos immunoreactivity following a 1 mg/kg CNO injection followed by a 10 mg/kg cocaine prime. The number of c-Fos+ cells in the LA was lower in the Gi-DREADD group that received CNO compared to those expressing the control virus that received CNO (t(6)=2.47, p=0.049), indicating that CNO in Gi-DREADD rats reduced the number of active cells following cocaine exposure (Supplemental Figure 8A).

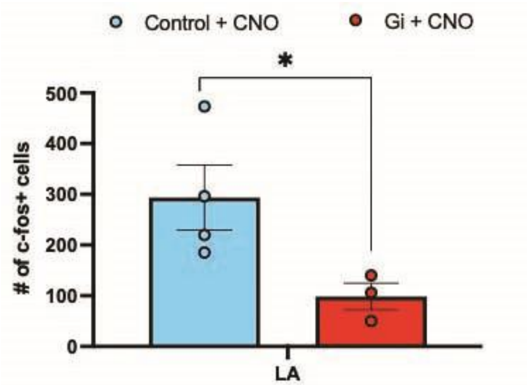


**Supplemental Figure 8. CNO administration in Gi-DREADD rats impacts c-Fos immunoreactivity.** Rats (n=7, Control + CNO=4, Gi + CNO=3) expressing the control virus and rats expressing the Gi-DREADD virus were given i.p. injections of 1 mg/kg CNO before a 10 mg/kg cocaine prime injection. c-Fos immunoreactivity in the LA was examined and the number of c-Fos+ cells in LA was lower in Gi-DREADD animals relative to controls **(A)**. Graph shows group means ± SEM. *p<0.05.

**Administration of CNO in LA of Gi-DREADD rats decreases c-Fos immunoreactivity after cue-induced reinstatement.**

To confirm that CNO affected neuronal activity in the LA of DREADD expressing rats, we examined c-Fos immunoreactivity following a cue-induced reinstatement session since c-Fos expression serves as an indicator of activity. CNO or ACSF microinfusions were administered into the LA of Gi-DREADD expressing, Gq-DREADD expressing and control rats prior to the session. Analysis of c-Fos+ cells yielded a main effect of treatment ($F_{(2,13)}$=5.78, p=0.016). Gi-DREADD rats that received a CNO microinfusion had fewer c-Fos+ cells than DREADD controls (p=0.039) and Gq-DREADD animals that received CNO (p=0.022), indicating that CNO administration reduced the number of active LA cells following cue-induced reinstatement (Supplemental Figure 9A). Behaviorally, there was no significant effect of treatment ($F_{(2,20)}$=0.77, p=0.48), but a trend for Gi-DREADD rats given CNO to make fewer active lever presses than Gq-DREADD rats given CNO (p=0.09, Supplemental Figure 9B).

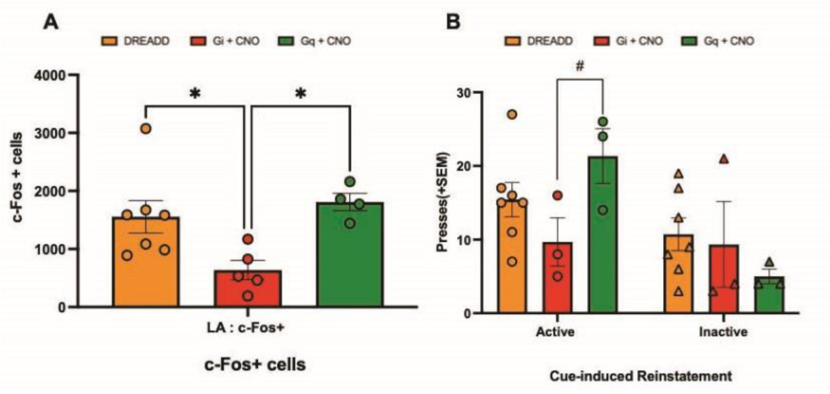


**Supplemental Figure 9. Administration of CNO in LA of Gi-DREADD rats diminishes c-Fos immunoreactivity after cue-induced reinstatement.** Rats (n=16, Control=7, Gi + CNO=5, Gq + CNO=4) underwent a cue-induced reinstatement session after LA microinfusion of CNO or ACSF. c-Fos immunoreactivity was then examined as an indicator of cell activity during the session. Gi-DREADD rats that received CNO microinfusions had fewer c-Fos+ cells compared to DREADD controls and Gq-DREADD rats that received CNO **(A)**. Gq-DREADD expressing rats that received CNO were trending to make more active presses during cue-induced reinstatement than Gi-DREADD expressing rats that were treated wth CNO **(B)**. Graphs show group means ± SEM. #p<0.1, *p<0.05
